# Pretreatment of aged mice with retinoic acid restores alveolar regeneration via upregulation of reciprocal PDGFRA signaling

**DOI:** 10.1101/2020.05.06.080994

**Authors:** Jason J. Gokey, John Snowball, Jenna Green, Marion Waltamath, Jillian J. Spinney, Katharine E. Black, Lida P. Hariri, Yan Xu, Anne-Karina T. Perl

**Author notes:** Corresponding author: Anne-Karina Perl. Grant support: R01 HL131661 (to JG, YX, JS, and AKTP), T32 HL007752 (to JJG), U01 HL122642 (YX and AKTP) LungMAP, U01 HL122638 (YX and AKTP) LungMAP, U01 HL134745 (YX and AKTP) PCTC, K08 HL133603 (KEB), K23 HL132120 (LPH), Translational Fibrosis Academic and Research Committee funding CCHMC.

## Abstract

**Objectives:** Idiopathic Pulmonary Fibrosis (IPF) primarily affects the aged population and is characterized by failure of alveolar regeneration leading to loss of alveolar type 1 cells (AT1). Aged mouse models of lung repair have demonstrated that regeneration fails with increased age. Mouse and rat lung repair models have shown retinoic acid (RA) treatment can restore alveolar regeneration. Herein we seek to determine the signaling mechanisms by which RA treatment prior to injury supports alveolar differentiation.

**Design:** Partial pneumonectomy (PNX) lung injury model and next generation sequencing of sorted cell populations are used to uncover molecular targets regulating alveolar repair. *In-vitro* organoids generated from Mouse or IPF patient epithelial cells co-cultured with young, aged, or RA pretreated murine mesenchyme are used to test potential targets.

**Main outcome measurements:** Known alveolar epithelial cell differentiation markers, including HOPX and AGER for AT1 cells are used to assess outcome of treatments.

**Results:** Gene expression analysis of sorted fibroblasts and epithelial cells isolated from lungs of young, aged, and RA treated aged mice predicted increased PDGFA signaling that coincided with regeneration and alveolar epithelial differentiation. Addition of PDGFA induced AT1 and AT2 alveolar differentiation in both mouse and human IPF lung organoids generated with aged fibroblasts and PDGFA monoclonal antibody blocked AT1 cell differentiation in organoids generated with young murine fibroblasts.

**Conclusions:** Our data support the concept that reciprocal PDGFA signaling activates regenerative fibroblasts that support alveolar epithelial cell differentiation and repair, providing a potential therapeutic strategy to influence the pathogenesis of IPF.

**Key Question:** Which epithelial-mesenchymal crosstalk pathways are activated by RA pretreatment of aged lungs that support realveolarization after partial pneumonectomy surgery?

**Bottom Line:** Increased PDGFA/PDGFRA signaling in aged lungs promotes regenerative activation of interstitial matrixfibroblast which is required for AT2 to AT1 differentiation and alveolar regeneration.

**Read On:** *In-vitro* and in-vivo analysis demonstrated that PDGFA signaling supports alveolar matrixfibroblast and AT1 epithelial cell differentiation, both necessary for alveolar regeneration in aged lungs.

## Introduction

Several lung diseases are associated with advanced age, including lung cancers, chronic obstructive pulmonary disorder (COPD), and Idiopathic Pulmonary Fibrosis (IPF). IPF is characterized by extensive fibrosis, hyperproliferation of abnormal alveolar type 2 cells (AT2) and loss of alveolar type 1 cells (AT1) causing progressive respiratory decline and mortality usually within 5 years of diagnosis [1-3]. While the pathogenesis of IPF remains unclear, chronic alveolar epithelial cell injury and fibrotic activation of fibroblasts are linked to the disorder [4]. However, an understanding of the epithelial-mesenchymal crosstalk during chronic epithelial injury and in the context of aging, remains limited.

Reciprocal communication between epithelial and mesenchymal cells drives lung branching morphogenesis and epithelial differentiation during lung development and repair [5-7]. Single cell RNA analyses have provided more clarity regarding the heterogeneity of normal epithelial and mesenchymal cells during development. Single cell seq analyses have identified several distinct heterogeneous pulmonary fibroblast populations [7, 8]. Of all pulmonary stromal cells, the mesenchymal alveolar niche cells (MANCs) [7] and LGR5 positive resident alveolar fibroblasts [9] have received recent attention due to their capacity to support the alveolar niche and differentiation of the alveolar epithelium. When used in alveolar organoid cultures MANCS support large organoid formation with high AT1/AT2 ratios. MANCs are Wnt responsive, AXIN2+ cells that also express PDGFRA and transcriptionally overlap with the *Lgr5* positive resident alveolar fibroblasts. In addition to the PDGFRA positive MANCS, there are WNT2+/PDGFRA+ and PDGFRA+ interstitial resident fibroblast populations [7]. An important role of these mixed PDGFRA expressing fibroblast populations in normal septation, realveolarization, and in bleomycin-induced fibrosis has been demonstrated by multiple studies [10-19]. Interstitial PDGFRA+ fibroblasts consist of mixed populations consisting of three distinct fibroblast populations with specific biological functions: contractile myofibroblasts, matrix synthesizing and remodeling matrixfibroblasts [10, 11, 14, 15] and lipid storing lipofibroblasts [20, 21]. Myofibroblasts, which express low levels of PDGFRA, induce smooth muscle actin expression during postnatal secondary septation and become activated during alveolar regeneration in young adult mice [14-17]. How the “activation” of lung interstitial fibroblasts occurs during regeneration remains unclear. In contrast to findings in young mice, aged mice fail to regenerate alveolar septate after PNX in association with decreased cell proliferation and increased differentiation of myofibroblast [22]. In other lung re-alveolarization models, retinoic acid reinitiates septation in young adult rats and mice [16, 23, 24]. In the present study, we demonstrate that preconditioning of aged mice with retinoic acid treatment before PNX can reinitiate alveolar septation. We used this combination of retinoic acid conditioning followed by PNX in aged mice to further study the epithelial-mesenchymal crosstalk that is important for “regenerative activation” of interstitial lung fibroblasts. These studies provide insights into future treatment options for lung regeneration and fibrosis resolution in IPF.

## Materials and Methods

### Mouse husbandry and partial left lobe pneumonectomy (PNX)

Wildtype mixed background mice were used for flow cytometry analyses and organoid culture. Young mice were 12-16 weeks of age and aged mice were >40 weeks. Aged RA pretreatment mice were treated with *trans*-RA (Sigma R2625) at a dose of 2ug/g BW dissolved in DMSO and peanut oil administered via intraperitoneal injection daily for 10 days prior to pneumonectomy with a 2 day break after the first 5 days. Mice were then subjected to PNX or sham surgery (Sham) and harvested 5 or 21 days post-surgery [25]. For each experimental group (Young Sham, Young PNX, Aged PNX, and Aged RA pretreated) N>3 were collected for each experiment.

### Patient samples

Donor patient samples were obtained from healthy lungs rejected for transplant and the Cincinnati Children’s Hospital Medical Center Institutional Review Board declared that donor tissue samples were Institutional Review exempt, in accordance with protocol 2013-3356. IPF patient samples were collected from transplanted lungs and informed consent was obtained from each subject in accordance with the Partners Institutional Review Board (2013P002332), Boston, Massachusetts, USA. Patient clinical data is available in Supplemental Table S6.

### SHG and immunofluorescence

IPF and donor fixed lung tissue in OCT blocks were sectioned at 250um and cleared via the PACT protocol followed by whole mount antibody stain. In short: OCT is removed with PBS and then tissue is placed in a cold 4% polyacrylamide (1610140 Bio-Rad) hydrogel with 0.25% photoinitiator (VA-044, Wako) solution overnight. The tissue was polymerized at 37° for 4 hours, washed with PBS to remove excess hydrogel and then incubated at 37° overnight in an 8% SDS in PBS solution for permeabilization. After washing with PBS, a 25% Quadrol (122262-1L, Sigma Aldrich) solution in PBS was added for 16 hours on a 37° rotator, after which the samples were washed in PBS and ready for immunostaining. For immunostaining the PACT cleared tissue sample was incubated with 4% donkey serum blocking solution at room temperature, then incubated with primary antibodies aSMA (Sigma A5228) and Abca3 (Seven Hills WMAB-17G524) at 4° degrees for 3 days. Unbound antibody was removed with PBS followed by fluorescent labeling with secondary antibodies. After washing with PBS, the tissue was mounted in RIMS solution for second harmonic imaging on a Nikon FN I upright microscope and analyzed using Nikon Elements and Imaris software.

### Cell sorting and FACS

Cell sorting was performed using Miltenyi Biotec microbead positive selection. Human CD326+ or Mouse CD326+ antibody conjugated beads were used to isolate epithelial cells and Human or Mouse CD140+ conjugated beads were used to isolate PDGFRA+ fibroblasts. Isolation was performed as per manufacturer’s instructions using LS columns for positive selection. Flow Cytometry was performed using fluorescent labeled antibodies (Supplemental Table S7). Cells were sorted using a Becton Dickenson LSRII analyzer with 5 lasers.

### RNA analysis (qPCR and RNAseq)

RNA-Sequencing was done by CCHMC’s Gene Expression Core. The resulting fastq files were aligned to mm10 or GRCh37 using Bowtie2 (1). Raw gene counts were obtained using Bioconductor’s Genomic Alignment and normalized FPKM values were generated using Cufflinks (2,4). Mouse RNA-seq was performed on 12-16 week old and >12 month old mice following SHAM surgery or PNX surgery with or without RA pretreatment (n=2-3). Whole mouse lungs were dissociated using dispase (Corning) and sorted with CD140 or CD326 microbeads (Miltenyi Biotec) prior to sequencing. Human donor and IPF samples were also dissociated and microbead sorted similar to mouse samples for CD140+ prior to sequencing (n=3). DeSeq was used to analyze raw gene counts and calculate differentially expressed genes (3). To identify differentially expressed genes in mice, a cutoff of FC>2 and pvalue<.01 was used and a cut off of FC>1.5 and Pvalue<.05 was used for human samples. All differentially expressed genes were expressed with a FPKM>1 in at least half of the replicates in one of the conditions being compared.

Mouse RNA-seq gene expression patterns were determined by analyzing all genes differentially expressed after aged PNX or after aged RA pretreatment PNX when compared to aged SHAM. These differentially expressed genes were used to make a z-score normalized FPKM expression matrix using all samples. Partek Genomics Suite’s hierarchal clustering coupled with heatmap generation was then used to cluster and visualize expression patterns. ToppGene’s ToppFun was used to identify functional enrichment hits of significantly altered RNAs within particular patterns (5). Upstream driver analyses were performed using differentially expressed genes from a given comparisons using Ingenuity Pathway Analysis’s upstream analyses software. RT-PCR was performed by generating cDNA using iScript (Bio-Rad). TaqMan assay probes (Thermofisher) were used to assess RNA expression and probes used are tabulated in Supplemental Table S8.

### Organoid generation

Human and mouse lung tissue were dissociated into single cell suspensions using dispase and DNase. Cell suspensions were incubated with FcR blocking reagent in MACS buffer (Miltenyi Biotec) followed by isolation of CD140+ and CD326+ cells as described earlier. CD140+ fibroblasts were co-cultured with CD326^+^ cells in a ratio of 10:1. Mixed cells were combined with matrigel (laminin, coll-iv, entactin) in a 1:1 ratio and cultured on a transwell insert in 24-well plates and incubated at liquid-air interface in MTEC plus media (DMEM-Ham’s F-12, HEPES, penicillin & streptomycin, fungizone, insulin, transferrin, cholera toxin, EGF and bovine pituitary extract). Organoid cultures were grown for 21 days, then fixed for immunofluorescence analysis.

## Results

To identify the role of resident PDGFRA+ interstitial fibroblasts in repair of the aged lung we used the murine partial pneumonectomy (PNX) model and compared alveolar regeneration in 12-16 week old mice (hereafter referred to as “young mice”) to 9-12 month old (hereafter referred to as “aged mice”) [22]. Consistent with previously published data, regeneration in aged mice is decreased and incomplete by 21 days post-surgery [22, 26] (Figure 1A). Since retinoic acid reinitiates septation in other lung regeneration models [16, 23, 24], we hypothesized that treatment of aged mice with retinoic acid prior to PNX injury (hereafter referred to as “RA pretreatment”), would promote alveolar regeneration in aged lungs. Male and female aged mice were treated with RA for 10 days prior to PNX surgery. Lungs from these mice were analyzed 5 and 21 days post-surgery using histology and flowcytometry. RA pretreatment restored alveolar septation in aged mice (Figure 1A-B). We have previously shown that myofibroblast and matrixfibroblast orchestrate alveolar regeneration in young mice [15]. Myofibroblast were characterized by expression of CD140 (PDGFRA) and CD29, matrixfibroblast were characterized by expression of CD140 and CD34 [15]. Young mice actively regenerate their lung after PNX, with increased proliferation of CD326+ epithelial cells and CD140+/CD34+ matrixfibroblast. While epithelial proliferation (CD326) is still induced in aged mice after PNX, proliferation of CD140/CD34 matrixfibroblast is blocked. RA pretreatment of aged mice restored proliferation in CD140/CD34 matrixfibroblast (Figure 1C-E). These data suggest that matrixfibroblast proliferation plays an important role in alveolar repair.

**Figure 1:**
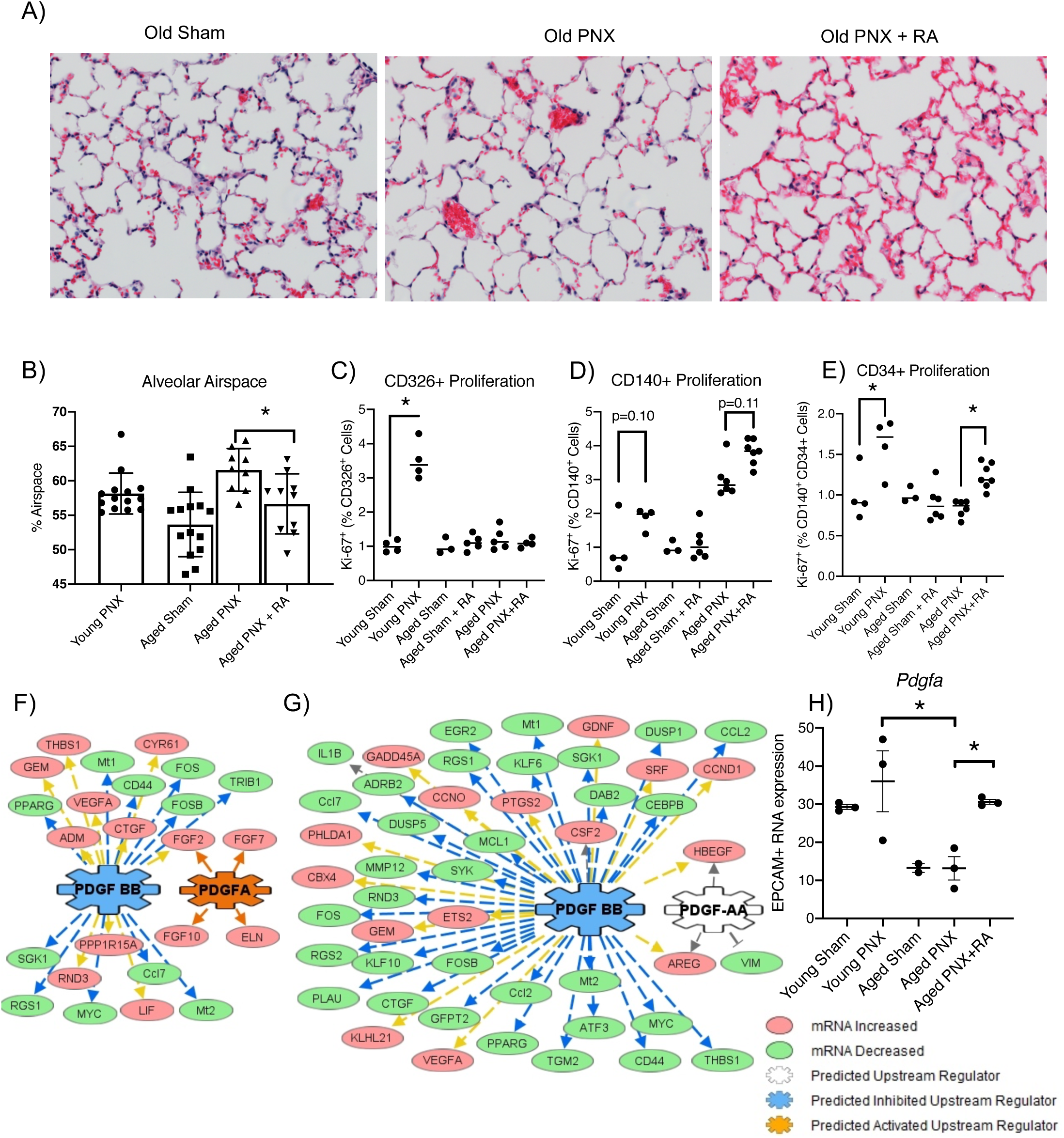
Mouse lung regeneration following RA pretreatment. A.) H&E stain of aged mice following sham surgery, Pneumonectomy (PNX) surgery, and RA pretreatment prior to PNX surgery. B.) Quantification of alveolar space and KI67 expression normalized to respective sham control in C.) CD326+, D.) CD140+, and E.) CD140+CD34+ cells in young and aged mice following SHAM, PNX and PNX after RA pretreatment. F,G.) Upstream analyses of RNA-Seq gene expression changes in PDGFRA+ mesenchymal cells from aged PNX mice when compared to either young PNX mice or aged mice pretreated with RA. In both comparisons, genes downstream of PDGF signaling were significantly altered. F) In PDGFRA+ mesenchymal cells isolated from aged PNX mice pretreated with RA, PDGF BB signaling was predicted to be significantly inhibited while PDGFA signaling was activated. G.) Similarly, in the aged PNX compared to young PNX, PDGF BB signaling was predicted to be significantly inhibited with PDGFA signaling being altered in the young PNX PDGFRA+ mesenchymal cells. H.) *Pdgfa* RNA expression in epithelial cells isolated from young sham injury, young PNX, aged sham injury, aged PNX, and RA pretreated aged PNX mouse lungs demonstrated increased *Pdgfa* expression in young PNX and RA pretreated aged PNX lung epithelial cells. * indicated p<0.05 as determined by one-way ANOVA followed by Tukey’s multiple comparisons test.

RNA sequencing of CD140+ mesenchymal and EPCAM+ epithelial cells isolated from sham, young, aged, and RA pretreatment mice were used to identify changes in gene expression associated with alveolar regeneration 5 days post PNX surgery (Supplemental Table 1, 2). Changes in gene expression patterns between PDGFRA+ fibroblasts and epithelial cells were strikingly different. In aged PDGFRA+ fibroblasts, a large subset of genes are altered by PNX while RA pretreatment prevented these changes (Supplemental Figure 1A). In contrast, in the epithelial cells, RA pretreatment induced a large subset of genes that were not changed by PNX (Supplemental Figure 1B). These data suggest that PNX in aged lungs induces “misdirected” gene activation and inactivation in PDGFRA+ fibroblasts that result in failure to regenerate. RA prevents these “misdirected” gene changes, allowing regeneration. In the epithelium of aged lungs, PNX does not induce gene changes, but RA pretreatment promotes gene changes that result in regeneration.

Functional enrichment analysis of the “misdirected” gene changes in PDGFRA+ fibroblasts show that RA pretreatment prevented the induction of genes associated with apoptosis, chromatin remodeling and inflammation, while simultaneously preventing the loss of genes associated with matrix organization, mesenchymal cell differentiation, lung development, stem cell differentiation, and regulation of epithelial cell proliferation (Supplemental Figure 1C). In contrast, functional enrichment analysis of the gene changes in epithelial cells shows that RA pretreatment inhibited expression of genes associated with inflammation and cytokine production while genes associated with epithelial development, branching, proliferation and differentiation were induced (Supplemental Figure 1D). These data demonstrate that in both PDGFRA+ fibroblasts and epithelial cells, inflammation suppression coupled with induction of cell type specific differentiation is required to allow for alveolar regeneration.

To identify gene networks associated with alveolar regeneration in PDGFRA+ fibroblasts, upstream driver analysis was performed using the gene changes observed between RA pretreatment and aged PDGFRA+ fibroblasts (Supplemental Table 3). Active PDGFA signaling and inactive PDGF BB signaling were predicted as important regulators of the gene changes that allow realveolarization (Figure 1F). Similarly, in young mice compared to aged mice (Supplemental Table 4), inactivation of PDGF BB signaling and alteration of PDGF AA were predicted to be primary mediators of the alveolar regeneration process (Figure 1G). Both PDGFA and PDGFB are expressed in the respiratory epithelium, suggesting paracrine signaling from the epithelium to the resident interstitial fibroblasts. RT-PCR was used to assess levels of *Pdgfa* in epithelial cells isolated from sham, young, aged, and aged RA pretreatment lungs. Epithelial *Pdgfa* was increased in young and RA pretreatment mice (Figure 1H). RNA analysis of *Pdgfb* demonstrated decreased *Pdgfb* signaling in the epithelium isolated from aged RA pretreatment lungs compared to aged lungs (Supplemental Figure 1E). Taken together these data suggest that increased reciprocal PDGFA signaling is essential for alveolar regeneration after PNX.

To test the role of PDGFA signaling in supporting alveolar regeneration and epithelial differentiation, an epithelial-mesenchymal mixed cell organoid system was utilized. Lung organoids were generated by combining primary EPCAM+ lung epithelial cells with primary PDGFRA+ fibroblasts isolated from young, aged, and RA pretreated murine lungs. Alveolar epithelial cell differentiation was assessed by immunofluorescence staining with SFTPC (AT2 cells), HOPX and AGER (AT1 cells) and quantified by morphometry (Figure 2A-B). SFTPC, AGER, and HOPX expression in primary lung epithelial cells from aged mice was increased when co-cultured in organoids with young or RA pretreated mouse lung PDGFRA+ fibroblasts, compared to organoids co-cultured with PDGFRA+ fibroblasts from aged mice. Independent of age and RA pretreatment, epithelial cells showed no difference in AT2/AT1 differentiation when cultured with young PDGFRA+ fibroblasts. While epithelial cells from young and RA pretreated mice expressed higher levels of *Pdgfa* (Figure 1H), their ability to differentiate into AT2/AT1 was impaired when cultured with aged PDGFRA+fibroblasts. When organoid cultures from old PDGFRA+fibroblasts were supplemented with PDGFA ligand, epithelial AT2/AT1 differentiation was restored as expression of AT1 cell markers, HOPX and AGER was increased (Figure 2B). Taken together these data demonstrate that the “age” of PDGFRA+ cells, but not epithelial cells, are important to promote epithelial differentiation in organoid cultures.

**Figure 2:**
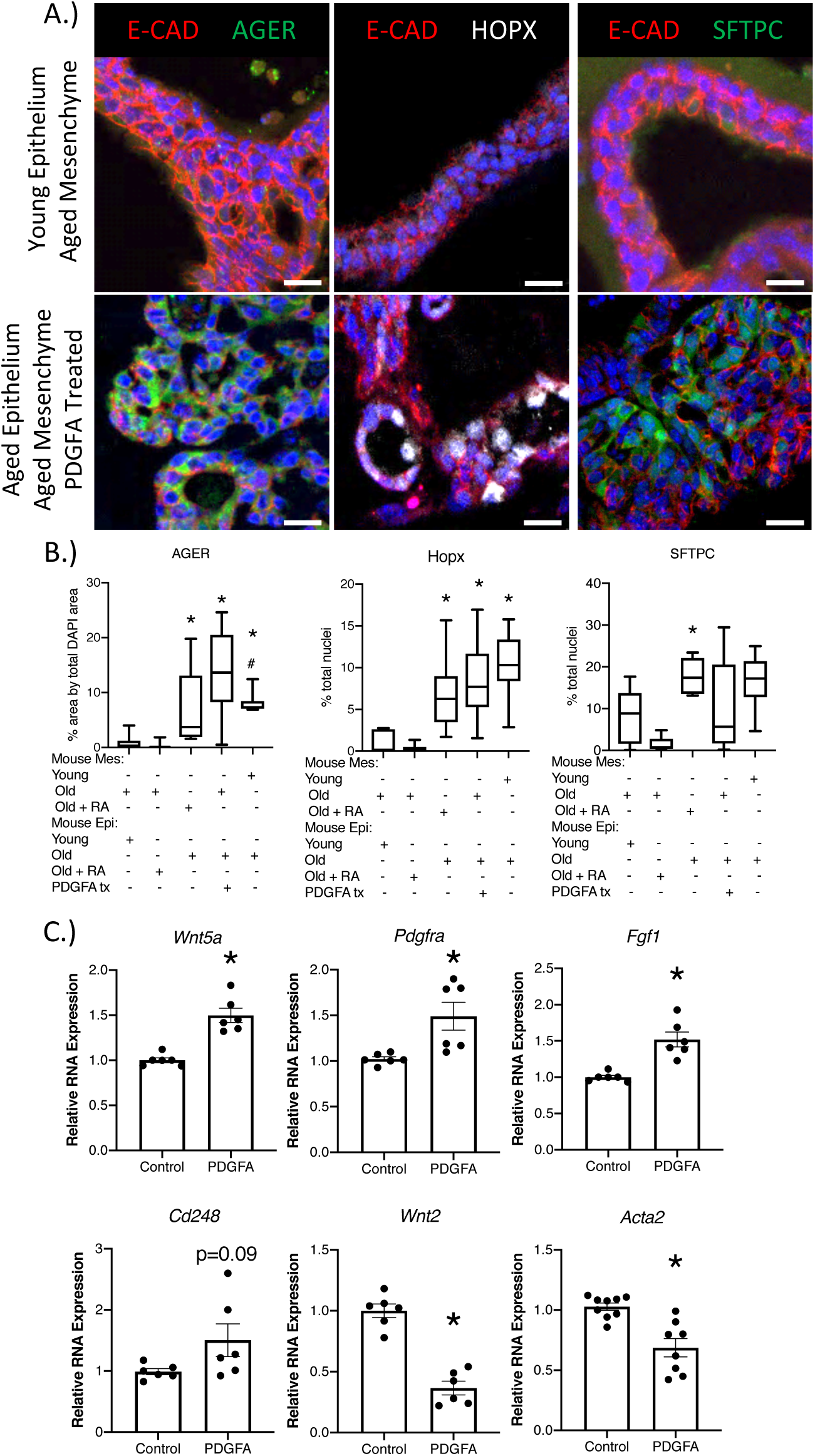
RA pretreatment and PDGFA treatment support alveolar differentiation in mouse lung organoids. A.) Immunofluorescence analysis of SFTPC, HOPX, and AGER in mouse lung organoids generated with young epithelium and aged PNX mesenchyme or aged epithelial and aged mesenchyme treated with PDGFA after 21 days in culture. B.) Quantification of total area masked by staining compared to total area of DAPI (AGER) or total cell counts as a percentage of total DAPI nuclei (HOPX and SFTPC) for the different combinations of mouse lung organoids generated demonstrates increased AGER and HOPX expression in organoids generated with young or old RA pretreated mesenchyme and PDGFA treated organoids generated with old mesenchyme. SFTPC expression was increased in organoids generated with RA pretreated old mesenchyme. C.) Quantification of RNA expression changes from IMR90 cells treated with PDGFA ligand demonstrated increased matrixfibroblast differentiation. * significant difference (P<0.05) as determined by one-way ANOVA followed by post hoc Dunn’s multiple comparisons test.

IMR90, human fetal lung fibroblasts express *PDGFRA*. To determine the role of epithelial PDGFA paracrine signaling on myofibroblast and matrixfibroblast differentiation, IMR90 cells were treated with PDGFA ligand. PDGFA treatment increased expression *CD248, WNT5a, FGF1* and *PDGFRA*, previously identified matrixfibroblast signature genes and reduced expression of *WNT2* and *ACTA2*, previously identified myofibroblast signature genes [8, 14] (Figure 2C). These data support the concept that epithelial PDGFA signaling promotes matrixfibroblast differentiation, which in turn support epithelial AT2/AT1 differentiation.

IPF is associated with advanced age, failure of alveolar repair and impaired AT1 differentiation. Based on the role of PDGFRA+ fibroblast in AT1 differentiation, we therefore examined PDGF signaling in IPF. Single cell analyses of the IPF and donor epithelium identified subsets of epithelial cells that produce high levels of PDGFB and PDGFA, with PDGFB signaling only being detected in IPF epithelial cells [6]. Due to altered PDGF expression in IPF lungs we hypothesized that the interstitial lung fibroblast population would be altered in IPF lungs. To assess the fibroblast populations composition changes of IPF patients, peripheral tissue of age-matched donor lungs and non-fibrotic peripheral areas of IPF lungs were digested into single cell suspensions and subjected to flow cytometry. Flow cytometry revealed a loss of 90% of CD140+ matrixfibroblasts in IPF (Supplemental Figure 2A). RNA sequencing of CD140+ fibroblasts from non-fibrotic areas of IPF lungs revealed changes in gene expression related to “extracellular matrix organization”, “response to wounding”, and altered “chromatin assembly”, when compared to donor lung CD140+ fibroblasts (Supplemental Figure 2B,C, Table 5). Further, upstream regulator analysis on differentially expressed genes in IPF predicted activation of PDGF BB signaling and reduced PDGF AA signaling in IPF, which is consistent with the finding in aged PNX injured mouse lungs (Supplemental Figure 2D). Together these data suggest a loss of normal interstitial matrixfibroblast as an underlying pathological feature of IPF.

Recent proteomics analysis of young and aged murine lungs [27] described downregulation of collagen XIV, which integrates collagen bundels by binding to collagen I fibrils and decorin [28]. Changes in extracellular matrix composition and distribution have been previously suggested in aged and IPF lungs [29]. To assess whether the loss of matrixfibroblast in non-fibrotic areas of the IPF lungs also results in changes of the extracellular matrix we performed Second Harmonic Generation resonance imaging (SHG) to visualize collagen and immunofluorescent analysis of smooth muscle actin and AT2 cell distribution in alveolar regions of normal (N=3) and IPF (N=3) lungs (Figure 3A). Normal lungs showed an intricate collagen network spanning in the lung parenchyma. This collagen structure was lost in otherwise fairly “normal” regions of the lung in patients with IPF (Figure 3A). Normal regions were determined by H&E staining of adjacent sections (Data not shown). Increased collagen deposition and smooth muscle actin is readily detected in fibrotic regions of IPF lungs (Figure 3A). These data demonstrate that the loss of the matrixfibroblast in the IPF lung result in destruction of the collagen network, revealing an underlying pathological feature of IPF.

**Figure 3:**
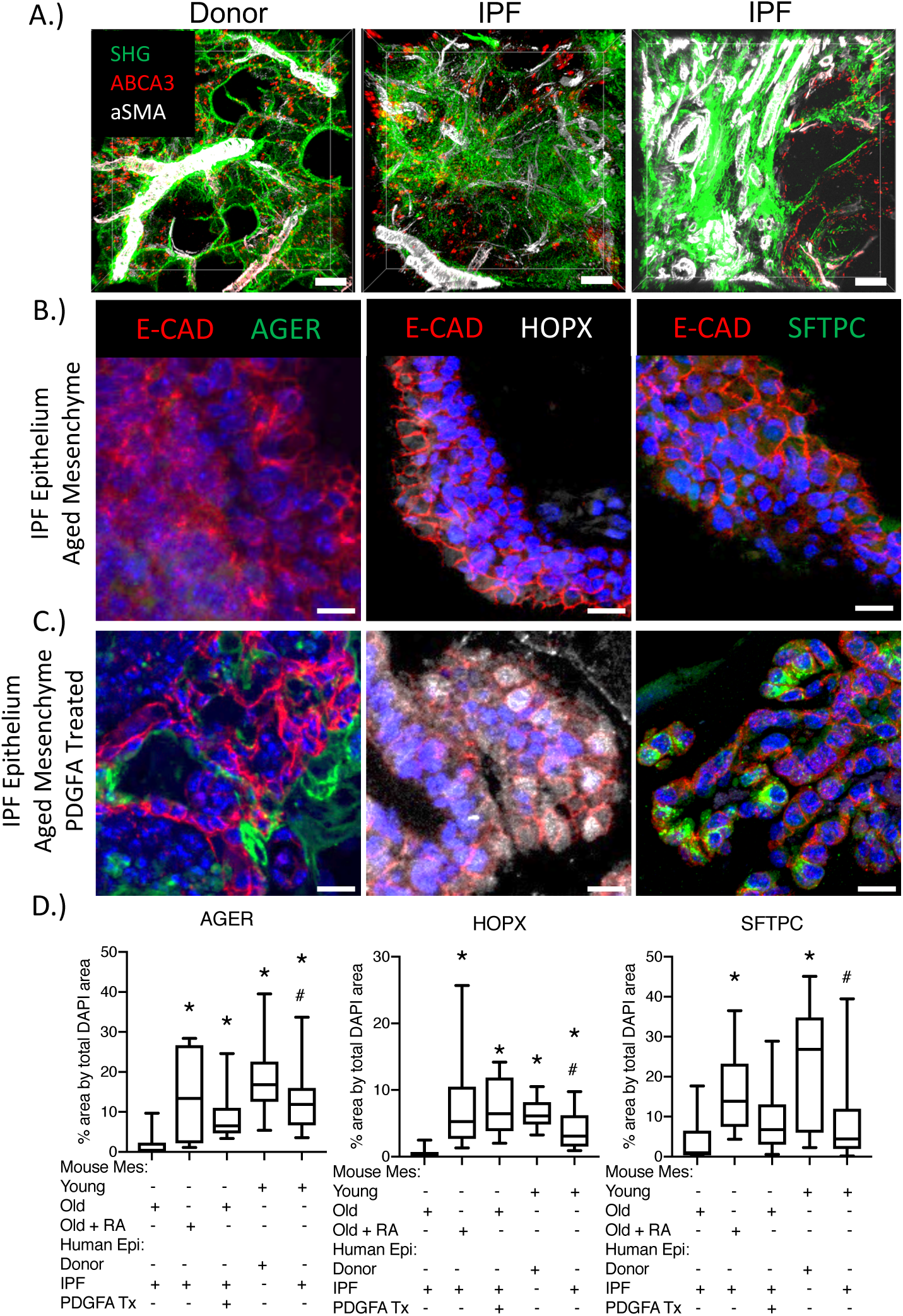
IPF lungs contain abnormal fibroblasts populations. A.) Second harmonic resonance imaging (green) showing structure of collagen in donor, “less severe” region of IPF, and “severe disease” region IPF lungs co-stained with immunofluorescence labeling of ABCA3+ AT2 cells (red) and aSMA+ smooth muscle (white), smooth muscle, in 250 μm thick lung sections. Scale bars indicate 50 μm. RA pretreatment and PDGFA treatment enhance expression of alveolar epithelial cell differentiation in human lung organoids. B-C.) Immunofluorescence analysis of AGER, HOPX, and SFTPC demonstrate increased expression in IPF epithelial cells cultured with aged mouse mesenchyme or treated with PDGFA. D.) Quantification of area imaged with AGER expression as normalized to total DAPI area. HOPX and SFTPC positive cells are normalized to total DAPI nuclei. * represents P<0.05 as determined by one-way ANOVA with post hoc Dunn’s multiple comparisons test compared to organoids generated with old mouse mesenchyme and human IPF epithelial cells.

IPF is characterized by the loss of AT1 cells, but it remains unclear if this is due to a defect of AT2 cells to differentiate and replenish chronically stressed AT1 cells [1]. Our data showed the loss of matrixfibroblasts in IPF and indicated that matrixfibroblasts are required for AT1 differentiation (Figure 3, Supplemental Figure 2). To test the hypothesis that loss of AT1 cells in IPF is due, at least in part, to loss of matrixfibroblasts, we assessed AT1 cell differentiation in human/mouse organoids. Organoids were generated by combining human CD326+ epithelial cells isolated from normal donor and IPF lungs with murine young, aged, or RA pretreated PDGFRA+ fibroblasts. We assessed AT2/AT1 differentiation in human donor or IPF epithelial cells with immunofluorescence analysis of HOPX, AGER, and SFTPC. Only young and RA treated fibroblasts supported AT1/AT2 differentiation in organoids in organoids generated from both donor and IPF epithelial cells (Figure 3 B,D). Other epithelial markers (P63, MUC5B, Acetylated Tubulin (ACTUB), and CCSP) were comparable among all combinations of fibroblast and epithelial cells; indicating there was not a shift in the bronchoalveolar/bronchial organoid ratio (Supplemental Figure 4). PDGFA treatment of organoids generated from old murine fibroblasts and human IPF epithelial cells enhanced AT1 (HOPX and AGER) and AT2 (SFTPC) cell differentiation to similar levels as organoids treated with RA or young fibroblasts (Figure 3C-D). These data suggest that epithelial cells from IPF patients have not lost the ability to differentiate into AT1 cells and that fibroblasts with active PDGFRA signaling is required for AT1 cell differentiation in IPF.

**Figure 4:**
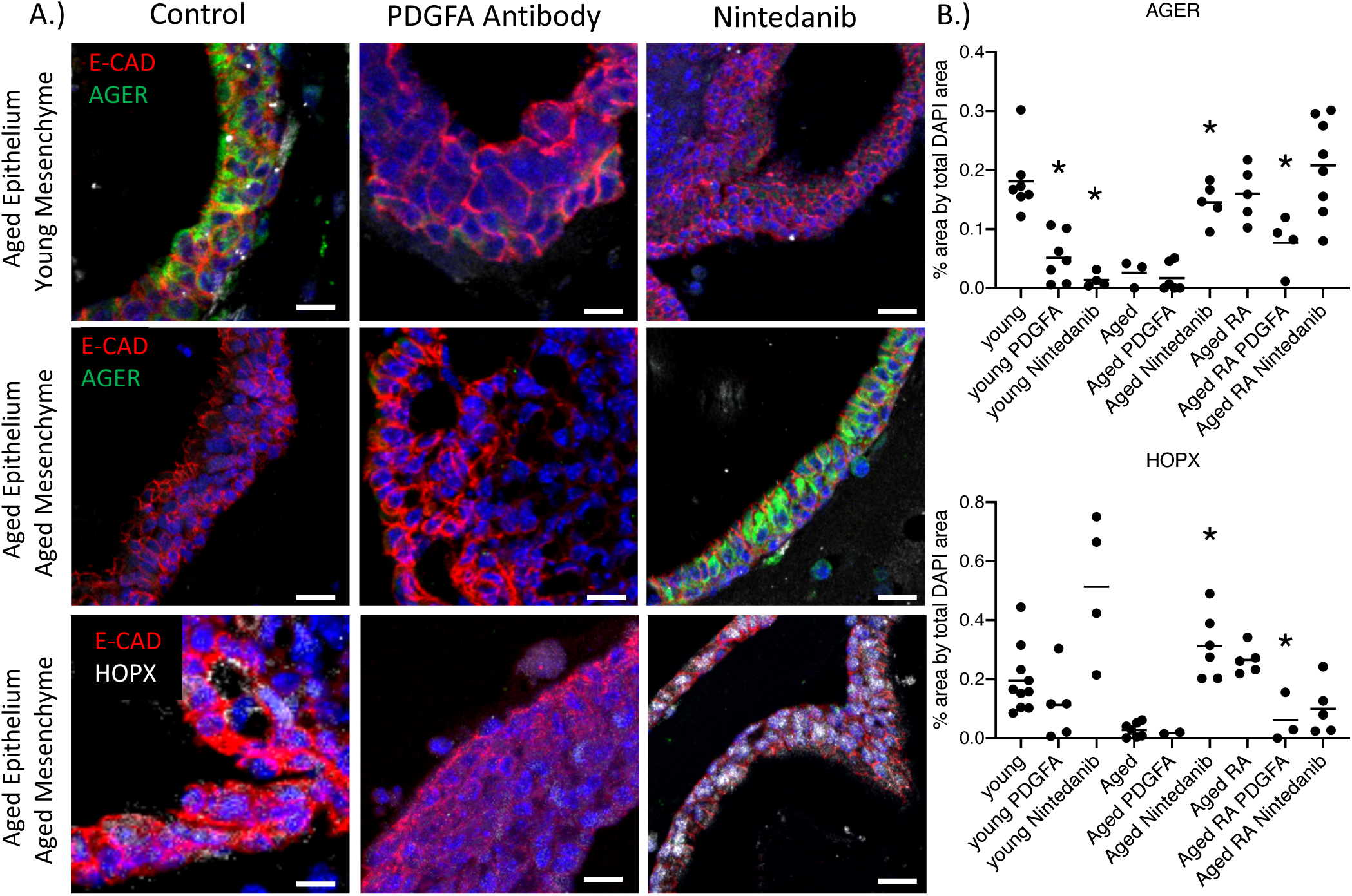
Inhibition of PDGFA blocks alveolar differentiation. Organoids generated with aged mouse epithelium co-cultured with young, aged, and aged RA treated mesenchyme. Organoids were treated with control, PDGFA neutralizing antibody, or nintedanib. A.) Immunofluorescence analysis demonstrates decreased expression of AGER and HOPX in organoids generated with young or RA treated aged mesenchyme following PDGFA antibody treatment. Nintedanib increased expression of AGER and HOPX in organoids generated with aged mesenchyme. B.) Quantification of AGER (normalized to total DAPI area) and HOPX (Normalized to total DAPI positive cell number) expression. * indicated p<0.05 as determined by ANOVA followed by Dunn’s multiple comparisons test.

To further test the importance of PDGFA signaling, aged mouse epithelial cells were co-cultured with young, aged, or RA pretreated PDGFRA+fibroblasts and treated with either PDGFA neutralizing antibody to directly inhibit PDGFA activity, or treated with nintedanib, a pan tyrosine kinase inhibitor, to block all PDGF (PDGFRA, PDGFRB and PDGFRC) receptors. PDGFA neutralizing antibody treatment reduced AGER, HOPX, and SFTPC expression in organoids generated with young and aged RA treated PDGFRA+fibroblasts (Figure 4A). Organoids generated with aged PDGFRA+fibroblasts had low expression of AGER and HOPX, however treatment with nintedanib enhanced expression of these AT1 markers (Figure 4B). These results demonstrate that specific inhibition of PDGFA ligand is sufficient to reduce AT1 differentiation. In contrast and consistent with previous publications, the pan tyrosine kinase inhibitor nintedanib increases AT1 cell differentiation in aged organoids, suggesting a ratio of PDGFA/PDGFB homeostasis is needed to support epithelial cell differentiation. However, in organoids, direct activation of PDGFA is sufficient to support alveolar differentiation, while inhibition of PDGFA is sufficient to block AT1 cell differentiation.

## Discussion

### Potential therapeutic advantage of PDGFA ligand treatment

The current focus on IPF research has revolved around correcting alveolar regeneration caused by the chronic epithelial cell injury which is believed to be an underlying cause of the disease. The FDA has approved two drugs, pirfenidone and nintedanib, a broad tyrosine kinase inhibitor that acts on several targets including PDGFRA and PDGFRB.

Thus far, these FDA approved treatments slow loss of Forced Vital Capacity but have not significantly enhanced life expectancy following diagnosis [30, 31]. Several recent clinical trials (phase 2 and 3) have continued along this path working to block other aberrant epithelial signaling pathways (mTOR, YAP, TGFβ) identified in IPF [5, 32]. However, perhaps a return to the fibrotic aspect of the disease would promote the identification of new targetable treatment options [33]. Fibroblasts support alveolar regeneration, and previous findings demonstrated RA pretreatment enhanced alveolar regeneration in animal models [23, 24], but RA treatment did not enhance alveolar repair in clinical trials treating COPD [34]. Likewise, direct treatment of mouse organoids with RA resulted in smaller organoid size with reduced epithelial cell differentiation, whereas inhibition of RA increased organoid size, increased epithelial cell proliferation via activation of YAP and FGF signaling [35]. These data suggest RA has a positive preconditioning effect but is inhibitory during active regeneration. In this study we used RA pretreatment prior to PNX injury to understand the downstream mechanisms that result in regenerative activation of fibroblasts leading to alveolar repair.

Our current findings support the notion that the activation of PDGFA signaling in RA treated fibroblasts lead to gene expression changes promoting epithelial regeneration. We also identified the epithelium of these mice as the compartment responsible for increased PDGFA ligand production. Bronchoalveolar organoids generated with aged mouse CD140+ fibroblasts fail to regenerate both AT2 and AT1 cells *in-vitro* regardless the types of epithelial cells used. However, treatment of these organoids with PDGFA ligand supports alveolar differentiation, suggesting PDGFA is sufficient to induce alveolar regeneration. PDGFA ligand appears to activate a cascade of epithelial-mesenchymal crosstalk, likely through enhanced mesenchymal differentiation into beneficial matrixfibroblasts, which promotes alveolar regeneration. Targeting downstream factors, such as PDGFA as potential therapeutic targets, may provide patients with IPF an efficacious treatment. Further study would be necessary to determine the efficacy of PDGFA ligand in models of interstitial lung diseases to exclude potential negative effects but targeting this signaling cascade may someday provide future therapies for IPF patients in need of efficacious treatment.

### The aging lung

Age is associated with increased risk for several diseases including IPF. Single cell RNA-seq analysis of aged murine lungs revealed altered transcriptional signaling that was associated with epigenetic changes in multiple cell types [27]. RA is known to place or remove epigenetic marks on histones and DNA [36]. In this study we demonstrate that RA pretreatment prevented many gene expression changes associated with aged fibroblasts and induced epithelial cell gene expression. These data suggest that RA pretreatment changed epigenetic landscaping of aged fibroblasts and allowed epithelial cells to respond to a regenerative stimulus.

It has been previously shown that monocyte recruitment is important for realveolarization after partial PNX [37]. Based on single cell RNA-seq analysis, aged mesenchymal cells have an increased pro-inflammatory signature [27]. Functional enrichment analysis in this study show that RA pretreatment prevented the induction of genes involved in inflammation in fibroblasts and inhibited their expression in epithelial cells. Taken together these data suggest that aged lungs are predispositioned to inflammation and that inflammation induced by PNX needs to be controlled to allow regeneration.

The Present study demonstrates that pulmonary fibroblast plays an important role for alveolar repair, which is lost with age, and highlights the ability of aged epithelium or epithelium from IPF patients to differentiate in the presence of regenerative matrixfibroblast. With age, PDGFA signaling is reduced in mouse epithelium. Direct supplementation of PDGFA In IPF human/mouse organoids can induce epithelial IPF AT1 cell differentiation. Upstream regulators analysis of disease related gene changes predicted increased PDGFBB and loss of PDGFA in IPF mesenchymal cells driving disease progression, consistent with previously observations [38]. In support of its putative role as a driver of IPF pathogenesis, PDGFBB increases fibrosis in other injury models, blocking PDGFBB reduces fibrosis in bleomycin-injured mouse lungs [39]. These findings, coupled with our currently described PDGFA neutralizing antibody and nintedanib treatment data, suggests that there is a balance between the fibrotic response induced by PDGFBB and the alveolar repair supported by PDGFA signaling. FDA approved nintedanib is a broad target tyrosine kinase inhibitor that blocks both PDGFA and PDGFB signaling cascades [30, 38, 39]. Our data suggest specific activation of PDGFA and specific inhibition of PDGFB may further enhance alveolar repair supporting a balance between PDGFA and PDGFB signaling cascades. Future therapies may rely on a combination of blocking aberrant pathways (PDGFRB), while supporting the activity of essential pathways for alveolar repair (PDGFA).

### Organoid model system

The present RNA analysis demonstrates epithelial and mesenchymal cell interactions is required for alveolar regeneration. Both human normal donor and IPF CD326+ epithelial cells have the potential to differentiate into AT2 and AT1 cells, suggesting the underlying mechanism for repair is still present in the IPF epithelium. This work suggests the mesenchyme activates PDGFA ligand in the epithelium to promote alveolar regeneration, a finding consistent with other reports showing the mesenchyme plays a key role in alveolar differentiation. Mesenchymal alveolar niche cells (MANCs) are a subset of WNT2+ AXIN2+ interstitial fibroblasts that interact with alveolar epithelial cells to support repair and development of alveolar region [7]. Another subset of interstitial fibroblasts, LGR5+ cells, also support alveolar differentiation [9]. These findings demonstrate the need to study alveolar regeneration and differentiation in a non-cell autonomous context. Herein, a relatively simple model of lung development and regeneration was utilized, in which interstitial fibroblasts and alveolar epithelial cells were co-cultured. This model demonstrates a need to address not only the aberrant alveolar epithelial cells in interstitial lung diseases such as IPF, but also take into consideration the fibroblasts, and potentially other cell types, that support alveolar remodeling during injury.

Taken together, our data suggest a model of paracrine signaling leading to alveolar repair in the aged lung, by which RA pretreatment of the mesenchyme induces signaling to the epithelium that activates epithelial PDGFA secretion. Production of PDGFA by alveolar epithelial cells then activates pulmonary matrixfibroblast differentiation, which in return leads to restoration of alveolar differentiation after injury.

## Figure Legends

**Supplemental Figure S1:** PDGFRA+ and EPCAM+ cells were analyzed separately. The effect of PNX on aged mice was determined by comparing aged Sham mice to aged PNX mice. PNX had little effect on the epithelial gene expression as seen by significant changes of only 166 genes but had a large effect on gene expression in PDGFRA+ mesenchymal cells; altering 2313 genes (Supplemental Table1). This pattern was reversed after RA pretreatment of aged mice. When compared to Sham, RA pretreatment of aged mice before PNX repressed many of the PDGFRA+ mesenchymal cells gene expression changes that occurred after PNX and altered the expression of a large subset of genes (2837) in the epithelial cell population that were not altered by PNX alone (Supplemental Table 2A, B). By combining the expression patterns of the genes that changed after PNX, with or without RA pretreatment, two large patterns emerged in both the PDGFRA+ mesenchymal population and EPCAM+ epithelial population C.) Functional enrichment of the genes in the indicated PDGFRA+ clusters shows RA pretreatment prevents the induction of genes associated with apoptosis, chromatin remolding, inflammation and immune responses while preventing the repression of genes associated with stem cell differentiation, mesenchymal cell differentiation, respiratory tube development, lung development, extracellular organization and regulation of epithelial cell proliferation. D). In the epithelium, following PNX, RA pretreatment of aged mice down regulated genes associated with immune responses, defense responses, inflammation and cytokine production and induced genes associated with epithelial development, branching, proliferation and differentiation in the epithelium of aged mice after PNX. E.) Analysis of *Pdgfb* expression in epithelial cells isolated from aged or RA Pretreated mouse lungs demonstrates RA pretreatment reduces *Pdgfb* RNA expression. * indicates p<0.05 as determined by a two-way unpaired T-Test.

**Supplemental Figure S2** A.) PDGFRA+ FACS sorting of single cell suspensions isolated from normal donor and non-diseased area of IPF lungs. Donor lungs contain approximately 13% PDGFRA+ mesenchymal cells whereas non-fibrotic regions of an IPF lung consist of 1% PDGFRA+ cells. B.) qPCR analysis of fibroblast associated gene expression from CD140+ isolated mesenchymal cells isolated from donor (N=5) and IPF (N=5) lung samples. RNA-seq of IPF PDGFRA+ lung cells compared to donor PDGFRA+ lung cells were preformed (n=3, 189 upregulated and 125 down regulated genes). C.) Functional enrichment analyses of differentially expressed genes demonstrated changes in extracellular matrix organization and proliferation. D.) Upstream analyses of the genes differentially expressed in PDGFRA+ mesenchymal cells from IPF and control donors predicted PDGF BB signaling to be activated in IPF and PDGF AA signaling to be altered.

**Supplemental Figure S3:** Immunofluorescent analysis of proximal epithelial cell markers in organoids generated with different combination of young and aged mouse epithelial and mesenchymal cells. A.) Muc5b (goblet cells), B) Acetylated Tubulin (ACTUB) (ciliated cells), C.) Scgb1a1 (CCSP) (secretory cells), and TP63 (basal cells) are present in all organoids regardless of combination of cells used to generate the organoid.

**Supplemental Figure S4:** Immunofluorescence analysis of proximal cell markers in human-mouse mixed cell organoids. A.) PGFA expression is increased in organoids generated with RA pretreated aged mouse mesenchyme. Immunofluorescence analysis of proximal cell markers MUC5B (B), ACTUB (C), CCSP (D), and TP63 did not change depending on mesenchyme used to generate the organoids. Statistical significance was determined using a one-way ANOVA followed by a Dunn’s multiple comparisons test.

**Supplemental Table S1:** EPCAM+ and PDGFRA+ sorted RNA-seq differentially expressed genes from aged Sham and aged PNX mice. Gene changes were determined using Deseq and differentially expressed genes were determined using a cutoff of FC>2 and pvalue<.01 was used.

**Supplemental Table S2:** EPCAM+ and PDGFRA+ sorted RNA-seq differentially expressed genes from aged Sham and aged RA pretreated after PNX mice. Gene changes were determined using Deseq to identify differentially expressed genes using a cutoff of FC>2 and pvalue<.01.

**Supplemental Table S3:** PDGFRA+ sorted RNA-seq differentially expressed genes from aged and RA pretreated after PNX. Gene changes were determined using Deseq and a cutoff of FC>2 and pvalue<.01 to identify differentially expressed genes.

**Supplemental Table S4:** PDGFRA+ sorted RNA-seq differentially expressed genes from young and aged mice after PNX. Differentially expressed genes determined using Deseq with a cutoff of FC>2 and pvalue<.01.

**Supplemental Table S5:** PDGFRA+ sorted RNA-seq differentially expressed genes from donor lung tissue and IPF lung tissue. Deseq was used to determine differentially expressed genes using cutoff of FC>1.5 and pvalue<.05.

**Supplemental Table S6:** Patient data. Tabulation of patient disease state, age, sex, history of smoking and race if information was disclosed.

**Supplemental Table S7:** Antibodies used, order information, and dilution used for immunofluorescence and FACs.

**Supplemental Table S8:** TaqMan qPCR probes used to assess RNA expression.

## Author Contributions

JJG, JS, JG, MW, JJS, KEB, LPH, YX, AKTP JJG, JS, AKTP conceptualized experiments; JJG, JG, MW, JJS performed experiments; JJG, JG, MW analyzed data; JS performed bioinformatic analysis; JJG, JS, KEB, LPH,YX, AKTP interpreted results; KEB, LPH provided clinical and pathological expertise; JJG, JS, AKTP wrote the manuscript and JG, MW, JJS, KEB, LPH, YX edited the manuscript.

## Acknowledgements

The authors thank the generosity of the patients who contribute to the advancement of science.

