## Supplementary figures and images for "Pretreatment of aged mice with retinoic acid restores alveolar regeneration via upregulation of reciprocal PDGFRA signaling"

### Supplemental Figure 1

Supplemental Figure 1

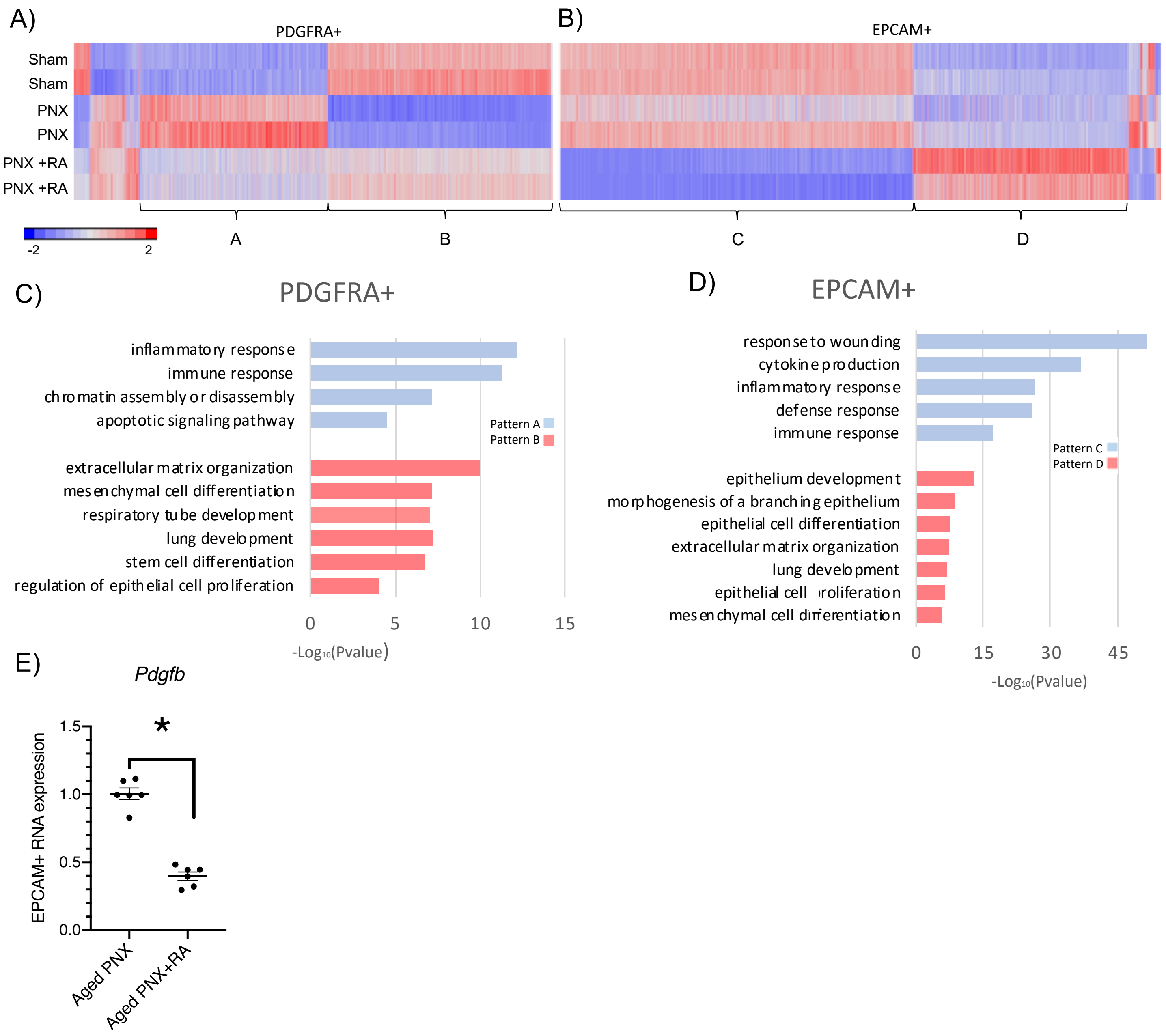

### Supplemental Figure 2

#### Supplemental Figure 2

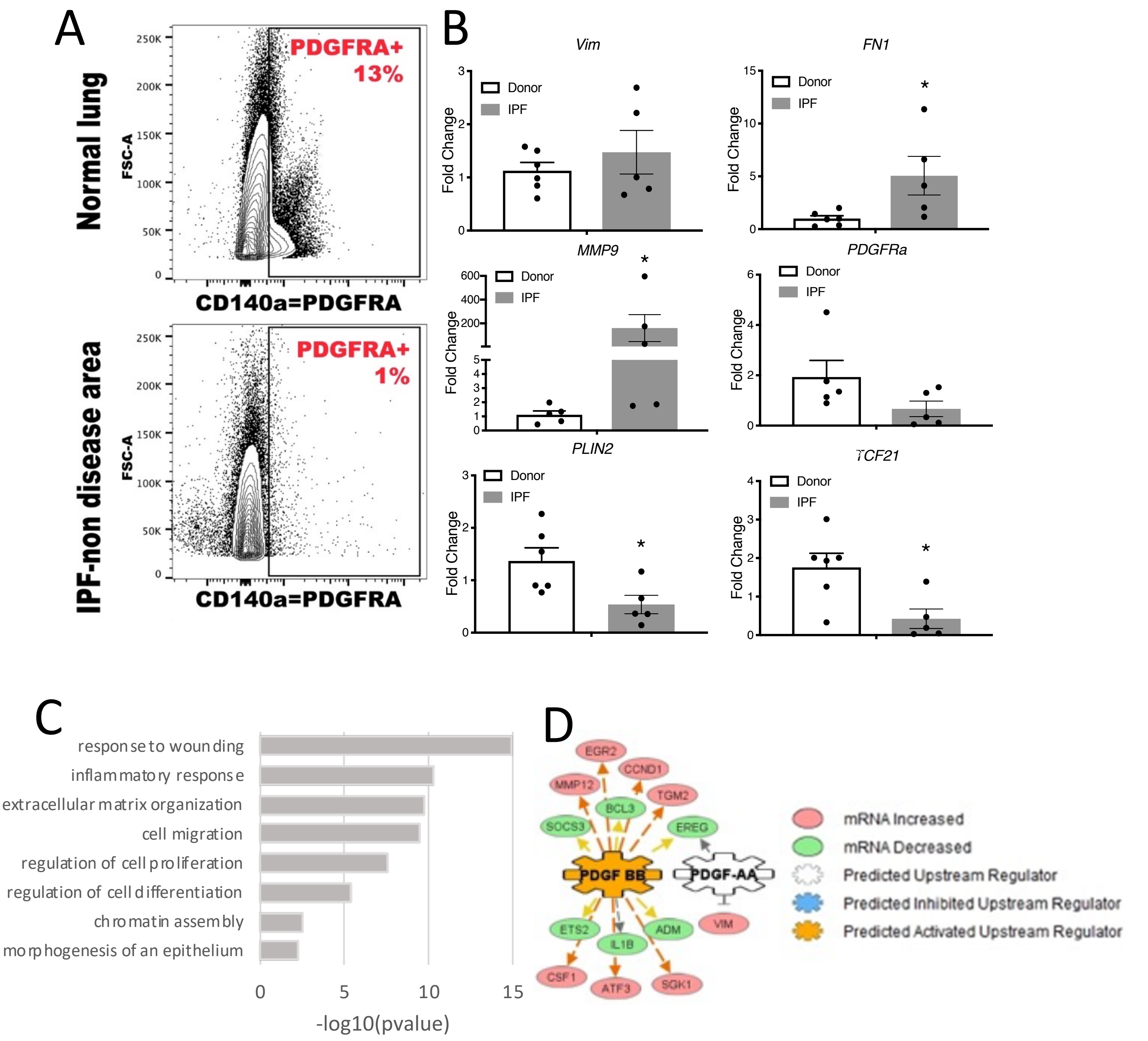

### Supplemental Figure 3

Supplemental Figure 3

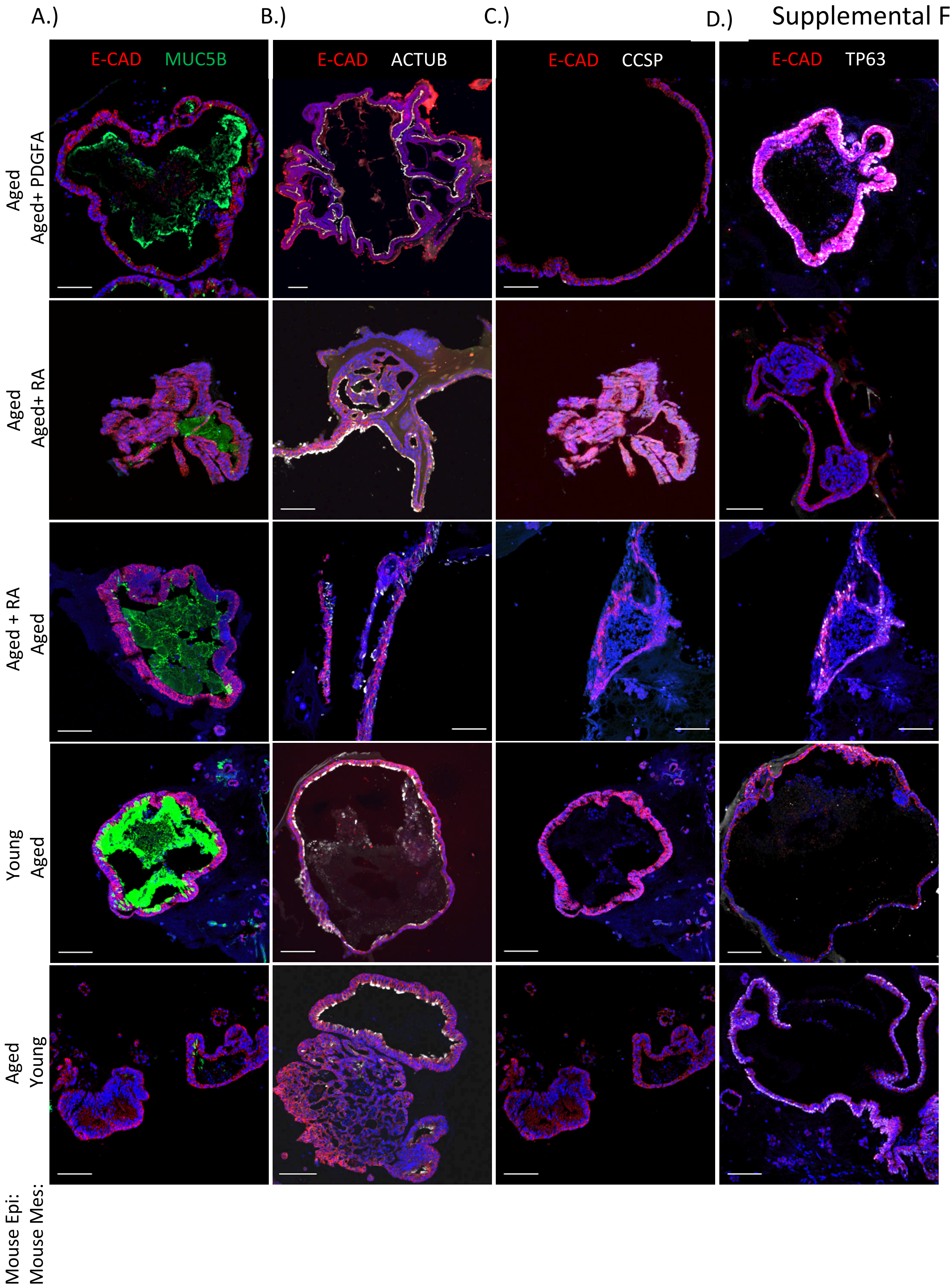

### Supplemental Figure 4

#### Supplemental Figure 4

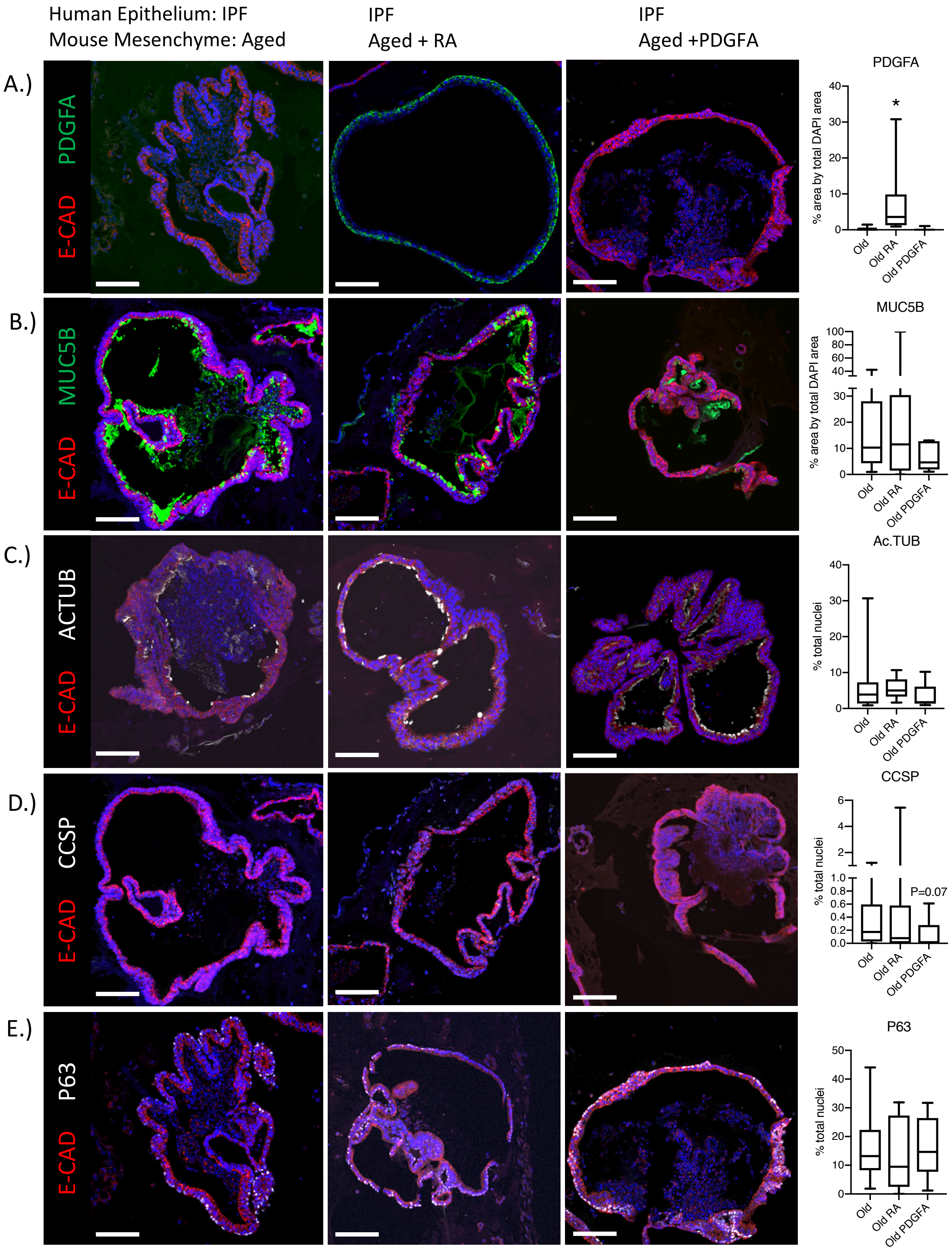
